# Vegetation monitoring using multispectral sensors – best practices and lessons learned from high latitudes

**DOI:** 10.1101/334730

**Authors:** Jakob J Assmann, Jeffrey T Kerby, Andrew M Cunliffe, Isla H Myers-Smith

## Abstract

Emerging drone technologies have the potential to revolutionise ecological monitoring. The rapid technological advances in recent years have dramatically increased affordability and ease of use of Unmanned Aerial Vehicles (UAVs) and associated sensors. Compact multispectral sensors, such as the Parrot Sequoia (Paris, France) and MicaSense RedEdge (Seattle WA, USA) capture spectrally accurate high-resolution (fine grain) imagery in visible and near-infrared parts of the electromagnetic spectrum, providing supplement to satellite and aircraft-based imagery. Observations of surface reflectance can be used to calculate vegetation indices such as the Normalised Difference Vegetation Index (NDVI) for productivity estimates and vegetation classification. Despite the advances in technology, challenges remain in capturing consistently high-quality data, particularly when operating in extreme environments such as the high latitudes. Here, we summarize three years of ecological monitoring with drone-based multispectral sensors in the remote Canadian Arctic. We discuss challenges, technical aspects and practical considerations, and highlight best practices that emerged from our experience, including: flight planning, factoring in weather conditions, and geolocation and radiometric calibration. We propose a standardised methodology based on established principles from remote sensing and our collective field experiences, using the Parrot Sequoia sensor as an example. With these good practises, multispectral sensors can provide meaningful spatial data that is reproducible and comparable across space and time.

## Introduction

Aerial imagery collected with drones is increasingly recognised by the ecological research community as an important tool for monitoring vegetation and ecosystems (Anderson and Gaston 2013, Salamí et al. 2014, Pádua et al. 2017, Torresan et al. 2017, Manfreda et al. 2018). Rapid advances in technology have resulted in increasing affordability and use of light-weight multispectral sensors for drones for a variety of scientific applications. Despite the increased presence of drone-sensor derived products in the published literature, standardized protocols and best practices for fine-grain, multispectral drone-based mapping have yet to be developed by the ecological research community (Manfreda et al. 2018). In this methods paper, we lay out the challenges of collecting and analysing multispectral data acquired with drone platforms and propose common protocols that could be implemented in the field, drawing from examples of applying drone technology to research in high-latitude ecosystems. The concepts developed herein are aimed at researchers with limited prior experience in remote sensing and spectroscopy, providing the tools and guidance needed to plan high quality drone-based multispectral data collection.

Multispectral imagery is widely used in satellite- and airplane-based remote sensing and has many benefits for vegetation monitoring when compared to conventional broad band visible-spectrum imagery. Including near-infrared parts of the spectrum, certain vegetation indices (VIs) can be calculated that allow for more detailed spectral discrimination among plant types and development stages. Such VIs can be highly useful for estimating biological parameters such as vegetation productivity and the leaf-area index (LAI; e.g. see Aasen et al. 2015, Wehrhan et al. 2016), and for the purpose of vegetation classification (Juszak et al. 2017, Ahmed et al. 2017, Müllerová et al. 2017, Samiappan et al. 2017, Dash et al. 2017). Particularly in remote high-latitude ecosystems, where satellite records suggest a ‘greening’ based on NDVI time series (Fraser et al. 2011, Guay et al. 2014, Ju and Masek 2016), multispectral drone monitoring could play an important role in validating satellite remotely-sensed productivity trends (see Laliberte et al. 2011, Matese et al. 2015).

A variety of multispectral camera and sensor options are available and have been deployed with drones. These range from modified off-the-shelf digital cameras (Lebourgeois et al. 2008, for examples see Berra et al. 2017, Müllerová et al. 2017), to compact purpose-build multi-band drone sensors such as the Parrot Sequoia (Ahmed et al. 2017, Fernández-Guisuraga et al. 2018) and the MicaSense Red-Edge (Samiappan et al. 2017, Dash et al. 2017). The Parrot Sequoia and MicaSense Red-Edge sensors are compact bundles (rigs) of 4-5 cameras with Complementary Metal-Oxide-Semiconductor (CMOS) (Weste 2011) sensors, a type of imaging sensor commonly found in the consumer cameras of phones and DSLRs. Each camera in the rig is equipped with an individual narrow-band filter that removes all but a discrete section of the visible and/or near-infrared parts of the spectrum (Table 1). New multispectral camera and sensor options continue to be released as technologies develop rapidly, yet many common considerations exist with the use of these type of sensors for the collection of vegetation monitoring data that we describe below.

**Table 1:**
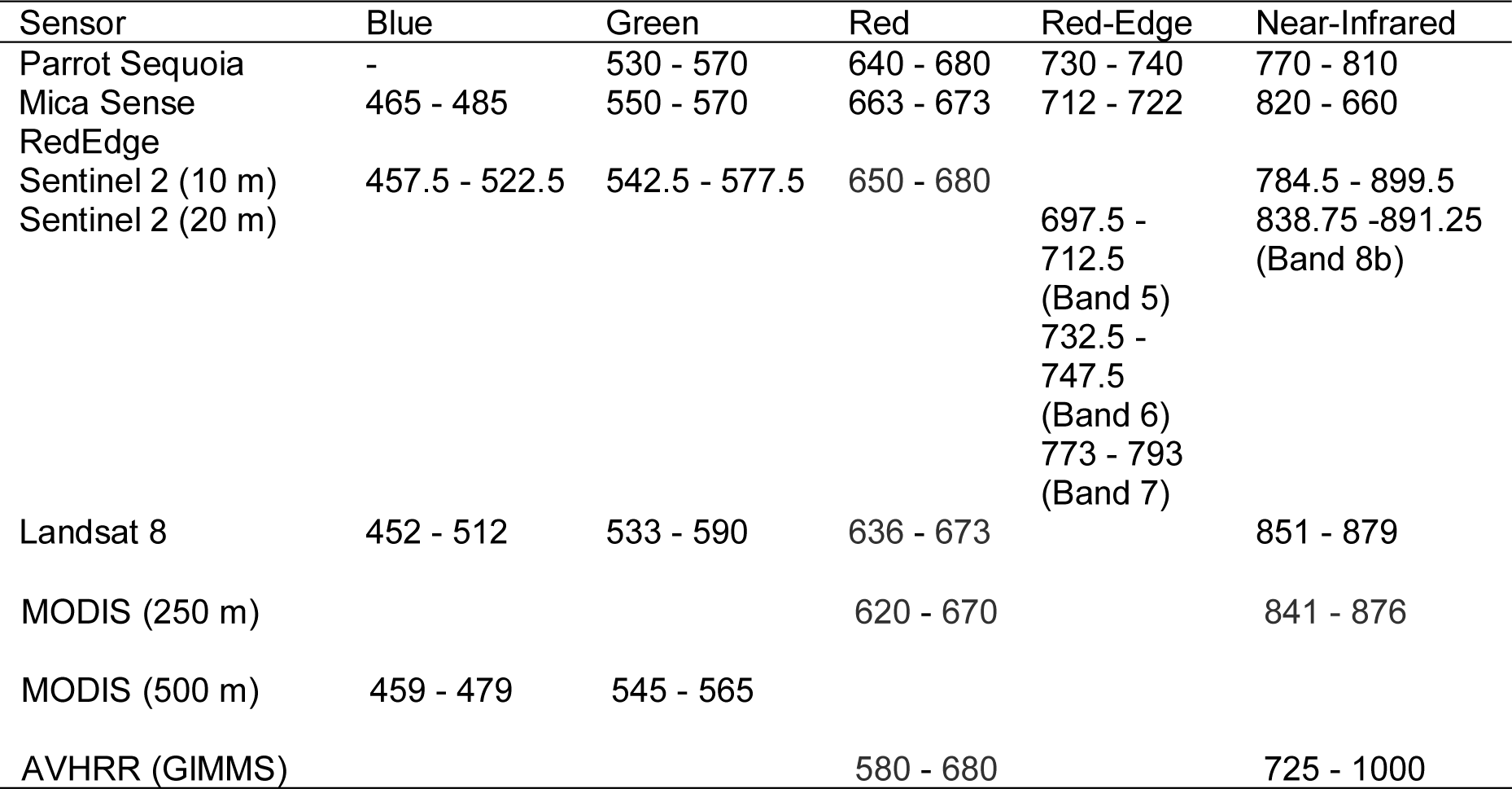
**Band wavelengths (nm) of the Parrot Sequoia and MicaSense Red-Edge Sensors with comparable Sentinel, Landsat, MODIS and AVHRR bands (Barnes et al. 1998, NOAA 2014, Barsi et al. 2014, European Space Agency 2015, MicaSense 2016a, 2016b). Vegetation indices such as the NDVI, derived from the read and near-infrared bands, can be notably affected by differences in spectral bandwidth. For the NDVI the position of the red band has been found to be of particular importance (Teillet 1997).**

The purpose-made design of the recent generation of multiband drone sensors provide many improvements that increase the ease of use, quality and accuracy of the collected multispectral aerial imagery. These include: precise co-registration of bands, characterised sensor responses, well defined narrow bands, sensor attitude correction, ambient light sensors, geo-tagged imagery, and seamless integration into photogrammetry software such as Pix4Dmapper (Pix4D SA, Lausanne, Switzerland) and PhotoScan Pro (Aigsoft, St. Petersburg, Russia). Despite these advances, acquiring multispectral drone imagery that is comparable across sensors, space, and time requires careful planning and best practices to minimise the effect of measurement errors caused by three main sources 1) differences among sensors and sensor units, 2) changes in ambient light (weather and position of sun), and 3) spatially-constraining the imagery (Kelcey and Lucieer 2012, Turner et al. 2014, Salamí et al. 2014, Aasen et al. 2015, Pádua et al. 2017).

With the goal of collecting comparable and reproducible drone imagery in mind, we discuss the fundamental technical background of multispectral drone sensors (Section 1), outline the proposed workflow for data collection and processing (Section 2) and conclude by reviewing the most important steps of the protocol in more detail (Section 3-6). These perspectives emerged from protocols originally developed for the High Latitude Drone Ecology Network (HiLDEN – arcticdrones.org), and build on examples drawn from data collected with a Parrot Sequoia at our focal study site Qikiqtaruk – Herschel Island (QHI), Yukon Territory, in north-western Canada and processed in Pix4Dmapper. Nonetheless, much of the discussed content should transfer directly to other multispectral drone sensors, including the MicaSense RedEge and Tetracam products, as well as to a lesser degree modified conventional cameras.

### Technical Background on Multispectral Drone Sensors (Section 1)

A fundamental aim of vegetation surveys with multispectral drone sensors is to measure surface reflectance across space for two or more specific bands of wavelengths (e.g. the red and near-infrared bands), which then serve as a base for calculating VIs (such as the NDVI) or to inform surface cover classifications. Reflectance is the fraction of incident light reflected at the interface of a surface. VIs enhance the characteristic electromagnetic reflectance signatures of different surfaces (such as bare ground, sparse or dense vegetation), whereas classifications often partition images based on these differences. Leaf structure and chlorophyll content influence the spectral signatures of plants, and VIs transform spectra-specific variability into single variables that can be related to other measures of vegetation productivity and leaf area index (LAI) (Tucker 1979, Guay et al. 2014, e.g. see Aasen et al. 2015). In practice, drone-based reflectance maps are usually created by collecting many overlapping images of an area of interest, which are then combined into a single orthomosaic (map) with a photogrammetry software package (such as Pix4Dmapper or Agisoft PhotoScan).

Reflectance is not directly measured by multispectral imaging sensors, instead they measure at-sensor radiance, the radiant flux received by the sensor (Figure 1). Surface reflectance is a property of the surface independent on the incident radiation (ambient light), whereas at-sensor radiance is a function of surface radiance (flux of radiation from the surface) and atmospheric disturbance between surface and sensor (see Wang and Myint 2015 for a detailed discussion). Surface radiance itself is highly dependent on the incident radiation, and disturbance between surface and sensors is often assumed to be negligible for drone-based surveys (Duffy et al. 2017). At-sensor radiance measurements are stored as arbitrary digital numbers (DN) in the image files for each band at a determined bit depth. Without modification, the DNs may serve as a proxy for relative differences of surface reflectance during the ambient light conditions of a particular survey, but if absolute surface reflectance measurements are desired - e.g. for cross site, sensor or time comparison - a conversion (“calibration”) of the digital numbers into absolute surface reflectance values is essential (Figure 1).

**Figure 1:**
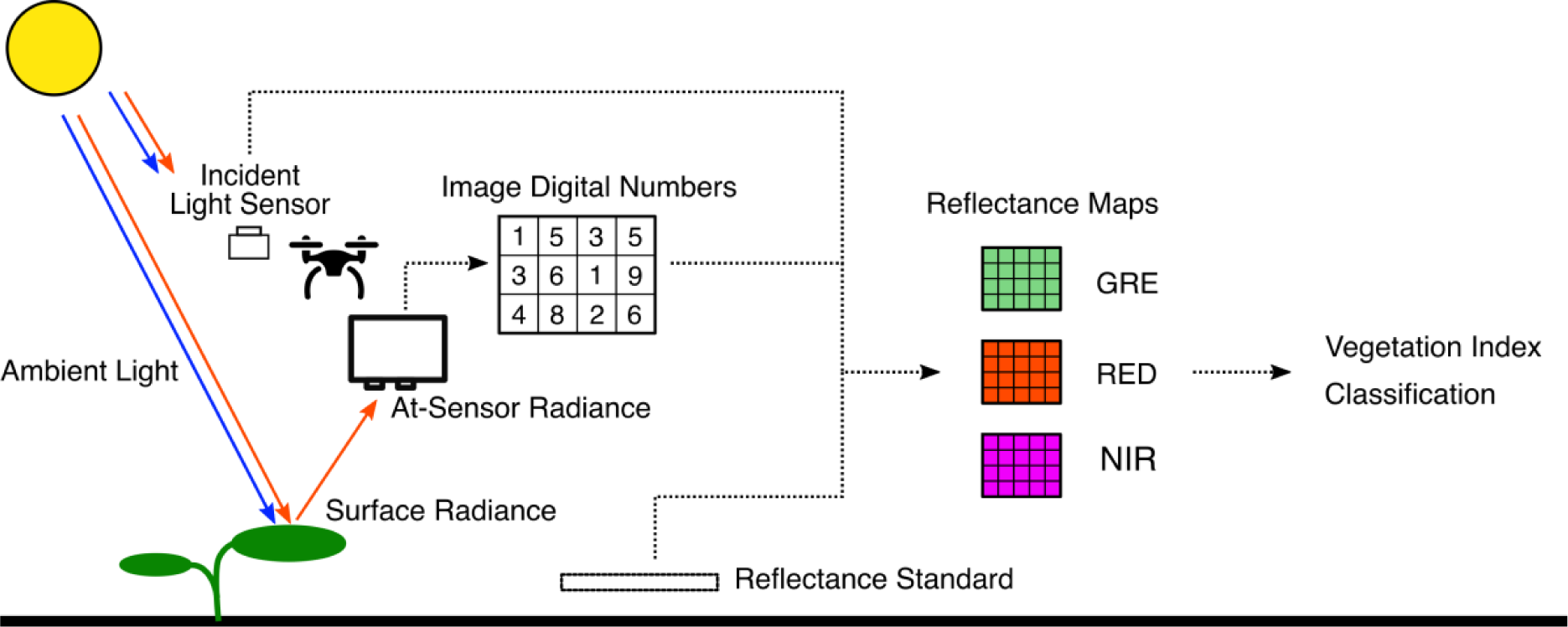
Simplified flow of information from surface radiance to reflectance maps using multispectral drone sensors. Surface radiance is measured as at-sensor radiance for each band by the drone sensor and saved as digital numbers (DNs) in an image file. Image DNs are then converted (“calibrated”) into reflectance values using an image of a reflectance standard acquired at the time point of the survey. The resulting reflectance maps for each of the sensor’s bands can then be used to calculate vegetation indices or as direct inputs for classification. Drone symbol by Mike Rowe from the Noun Project (CC-BY, http://thenounproject.com).

There are several ways to convert image DNs into absolute surface reflectance, but the most common is the so-called empirical line approach: Images of surfaces with known reflectance are used to establish an assumed linear relationship (empirical line) between image DNs and surface reflectance under the specific light conditions of the survey (Laliberte et al. 2011, Turner et al. 2014, Wang and Myint 2015, Aasen et al. 2015, Wehrhan et al. 2016, Ahmed et al. 2017, Crusiol et al. 2017, Dash et al. 2017). Additionally, information from incident light sensors, such as the Parrot Sequoia sunshine sensor may be incorporated to account for changes in irradiation during the flight. We would like to highlight here that this is not a calibration of the sensor itself, but a calibration of the output data. Practical aspects of radiometric calibration are discussed later in Section 6.

The relationship between DN and the surface reflectance value of a pixel is also influenced by the optical apparatus and the spectral response of the sensor, which require additional corrections (see Kelcey and Lucieer 2012 and, Wang and Myint 2015 for in-depth discussions). For the latest generation of sensors (e.g. MicaSense RedEdge and Parrot Sequoia) the processing software packages (such as Pix4Dmapper) automatically apply these corrections and little input is required from the user in this respect. Instructions on how to carry out the calibrations manually has been made available by some manufacturers (Parrot 2017a, MicaSense 2018c) and may be used by advanced users to develop their own processing workflow. However, understanding the principles of and why these corrections are required can be helpful to all users when planning multispectral drone surveys and handling the data outputs.

Firstly, the optical apparatus (i.e. filters and lenses) distort the light on its way to the sensor and therefore influence the relative amount of radiation reaching each pixel. Effects such as vignetting - pixels on the outsides of the images receive less light than those in the centre of the image (Kelcey and Lucieer 2012) – can produce desirable aesthetic effects in conventional photography, but bias data in different parts of the images when mapping surface reflectance. Converting the DNs of all pixels the same way would incorrectly estimate reflectance values towards the extremes of each image. This can be corrected for if the effects of the optical apparatus of the sensor have been characterised sufficiently (Kelcey and Lucieer 2012, Salamí et al. 2014).

Secondly, the relationship between DN and radiant flux is dependent on the sensitivity of the CMOS sensor unit in the specific band of the spectrum, the shutter speed, as well as the aperture and ISO value (signal current amplification at the sensor pixel level) settings during image capture. In the case of the Parrot Sequoia, this relationship is a linear function for which the parameters are characterised for each individual sensor unit at production. This is one of the major advantages of using purpose-built sensors such as the Parrot Sequoia and alike over modified consumer cameras. The relevant parameters of this relationship can be extracted from the image EXIF tags and applied to each image to obtain arbitrary reflectance values common to all Sequoias. These can then be converted into absolute reflectance using a standard of known reflectance (see Parrot 2017c).

When using Pix4Dmapper for processing Parrot Sequoia or MicaSense RedEdge data these corrections are automatically carried out by the software (Pix4d Personal Communication June 2017). Apart from defining the radiometric calibration image to establish the empirical line relationship, no additional input is required. The exact algorithms of Pix4Dmapper are proprietary and will likely remain a black box to the scientific community, and may change between software versions. To the best of our knowledge, at this time, there is no open source software currently available with the same scope and ease of handling of Pix4D mapper for processing multispectral drone data. During the completion of this manuscript, radiometric calibration features have been added to the recent release of Agisoft PhotoScan Pro (St. Petersburg, Russia).

#### Box 1

##### Quick Glossary

**Multispectral Drone Sensor**

A light-weight camera rig with at least two digital imaging sensors that capture monochromatic imagery in well-characterised and narrow bands of the electromagnetic spectrum. Often include bands outside the visible spectrum. Used to determine surface reflectance across space.

**Surface Reflectance**

Proportion of electromagnetic radiation reflected by a surface. Here specifically, the proportion of electromagnetic radiation reflected by a surface within narrow bands of the electromagnetic spectrum.

**Vegetation Index (VI)**

Mathematical transformation of surface reflectance values across multiple bands to allow for the estimation of vegetation productivity and surface cover type classifications.

**Digital Number (DN)**

Sensor-specific value used to denote strength of radiant flux to a sensor pixel. Arbitrary in nature, it requires knowledge of sensor response, optical apparatus and ambient light conditions to allow for conversion into surface reflectance values.

**Ground Sampling Distance (GSD)**

Distance between pixel centres or pixel-width measured on the ground of a digital aerial image.

**Ground Control Points (GCPs)**

Artificial or natural features with (often very accurately) known locations used to geo-rectify aerial imagery.

**Structure from Motion (SfM)**

Computational technique (computer vision) that uses relative positions of pixels from overlapping imagery of the same scene obtained at different angles to construct 3D models and composite orthomosaic images.

**Orthomosaic**

Mosaic of geometrically corrected (orthorectified) images so that scale is uniform across the mosaic from a nadir perspective (viewer 90° above viewing plane).

**Reflectance Map**

Orthomosaic of monochromatic imagery in a specific spectral band obtained with a multiband drone sensor. Pixel values contain (often radiometrically calibrated) surface reflectance values (ranging from 0 to 1). Can be used to calculate maps of vegetation indices.

### Data collection and processing – Workflow overview (Section 2)

Specific research questions and scientific objectives should be used to determine the exact methods used and the data outputs required from a multispectral drone survey (Figure 2). However, using a standardized workflow will help users avoid common pitfalls that affect data quality, and thus ensure repeatable and comparable data collection through time and across sites. We suggest starting by identifying the spatial and temporal scales required to address the research questions and scientific objectives (Step 1). Explicit consideration of scale is critical to the quantification and interpretation of any environmental pattern (Levin 1992), thus particular attention is required when planning drone surveys due to the scale-dependent nature of these inherently spatial data and its associated errors.

**Figure 2:**
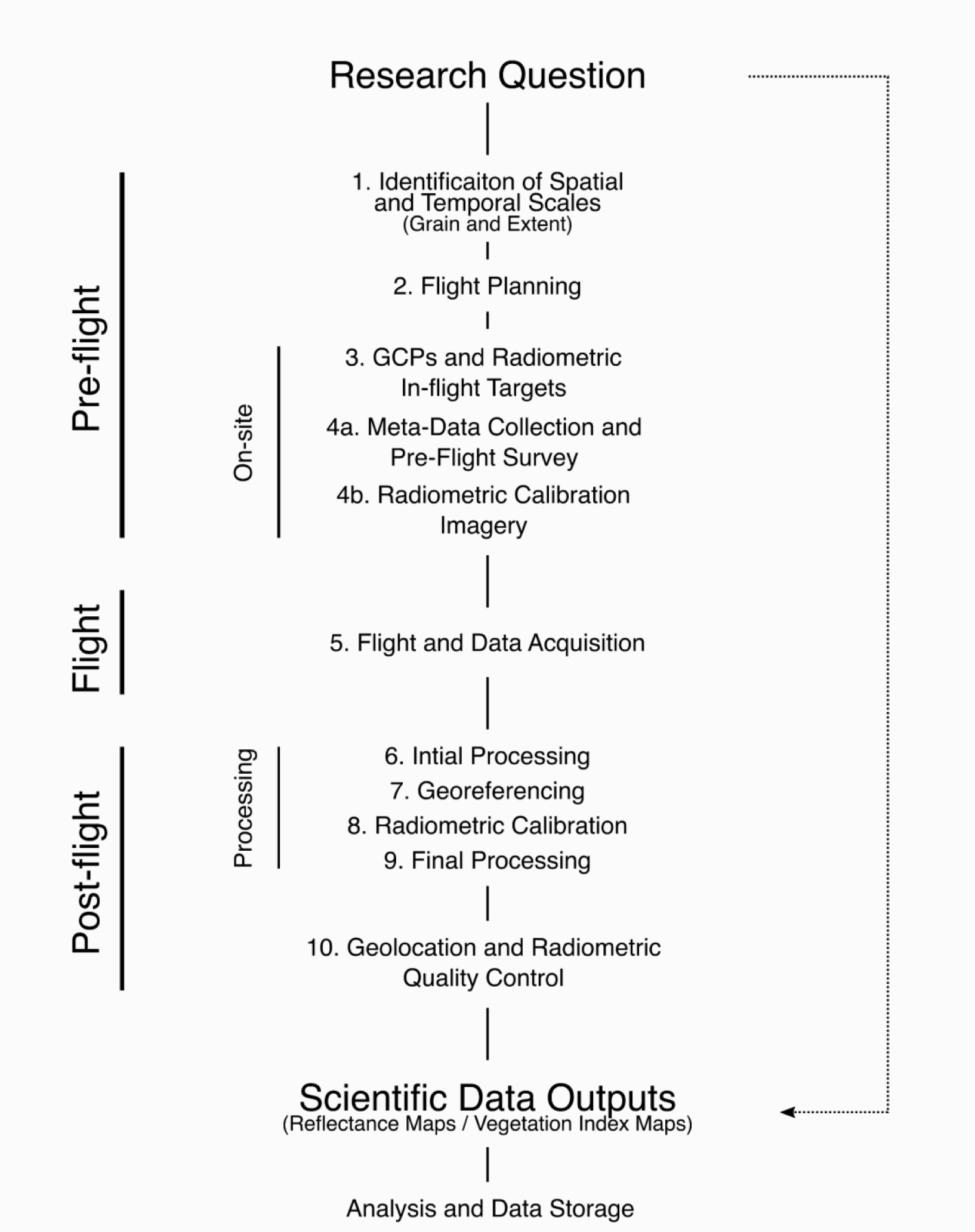
Overview of the proposed workflow for scientific data collection using multispectral drone sensors and guide to the sections of this publication. Flight planning is discussed in Sections 3 (Image Overlap and Ground Sampling Distance) and Section 4 (Weather and Sun) of this manuscript. Geo-location and use of ground control points (GCPs) in Section 5 and Radiometric Calibration in Section 6.

The selected spatial and temporal scales, together with the capabilities of the drone platform form the basis for flight planning (Step 2). Flight paths and image overlap (Section 3), as well as weather conditions and solar position (Section 4) are especially important to consider when planning multispectral drone surveys because of their impact on mosaicking and radiometric calibration. Once the flight plan is established, ground control points (GCPs) and radiometric in-flight targets need to be deployed on site, their locations determined with a high-accuracy global navigation satellite systems (GNSS) device (e.g. a survey-grade GPS receiver), and radiometric calibration imagery taken (Steps 3 and 4). We will discuss practical aspects of GCPs deployment and radiometric calibration in the final two sections (Section 5 and 6, respectively).

Once pre-flight preparations are completed, the drone is launched and the image data collected (Step 5). Though this may sound straight forward, in practice this can be challenging. Technical issues such as aircraft material failure, weather impacts on realized vs. planned flight path, and/or compass issues are not uncommon. Operator skill and logistical experience in the field should not be discounted, particularly when operating in extreme environments such as those found in the high latitudes (Duffy et al. 2017). Manufacturer guidance, online discussion boards and email lists (such as the HiLDEN network: arcticdrones.org) can provide help and information on these technical problems. Upon completion of the flight, image data can be retrieved from the sensors and transferred to a computer for processing. We recommend backing up the drone / sensor memory after every flight to reduce the risk of data loss due to hardware failure and crashes.

Processing will vary with the type of sensor / software that is used. Figure 2 outlines the core steps when processing Parrot Sequoia data with Pix4Dmapper Desktop. The initial processing step (Step 6) creates a rough model of the area surveyed using Structure from Motion – Multiview Stereo algorithms (SfM-MVS) (Westoby et al. 2012). The user then manually places GCP markers for geo-referencing (Step 7) and carries out the radiometric calibration (Step 8). These inputs are then incorporated by the software in a final processing step (Step 9), producing reflectance map and VI map outputs.

We suggest a final quality control step (Step 10) to assess the accuracy of the geo-location and radiometric calibration of the outputs, before using them in the analysis to answer the research questions. We also highlight that drone surveys can produce large amounts of data that can create challenges for data handling and archiving. It is helpful to produce a storage and archiving plan before data collection begins, test flights can provide valuable insights on data volume expectations for the project.

### Flight planning and overlap (Section 3)

A well-designed flight plan ensures that the full extent of the area of interest is covered at the appropriate grain size to fulfil the scientific objectives of the survey. The capabilities of drone and sensor, the terrain, as well as local regulations will constrain what is achievable. Flight planning software and manufacture guidance can assist and a wealth of information on flight planning and practise is available on the internet, including guidance on the legal aspects of operating drones in different jurisdictions. Here, we will focus on two aspects of mission planning particularly important for multispectral surveys: 1) image overlap - the proportion of overlap between neighbouring individual images in the pool of images covering the area of interest; and 2) spatial grain size or ground sampling distance (GSD) - the width of the ground area represented by each pixel in the imagery. Both are closely linked to, and limited by, flight height and speed, as well as sensor field of view and trigger rates.

Image overlap influences the percentage of pixels captured near to nadir view (sensor at 90° above surface of interest). Vegetative surfaces do not have lambertian reflectance properties; *i.e.*, they do not reflect light evenly in all directions, instead their reflectance is a function of both angle of incident light and angle of view. These relationships can be complex and are commonly described with so called bidirectional reflectance distribution functions (BRDFs) (Kimes 1983, for example Bicheron and Leroy 2000). For multispectral drone surveys non-uniform reflectance functions pose a challenge as they hamper the comparison of pixels captured at different angles of view (Aasen and Bolten 2018).

When obtaining surface reflectance imagery with wide-angled lenses, as those employed in many drone sensors, pixels near to the edges of the image have viewing angles notably different from 90° (up to 32° different for the horizontal field of view of the Parrot Sequoia). If a nadir angle of view (observer 90° above observed point) is assumed for these pixels the reflectance values in the extremes of the image maybe under or overestimated. High amounts of image overlap (75% - 90%) ensure that the whole area of interest is captured by pixels taken at near-nadir view. During processing these pixels can then be preferably selected as best estimates for surface reflectance at nadir view. Pix4D mapper carries out such a selection when creating reflectance maps (Pix4D Personal Communication, June 2017).

We recommend a minimum of 75% of for multispectral flights for both side- and front-lap (also recommended by MicaSense 2018a). However, more overlap might not always be better as there are penalties for very high amounts of overlap, affecting data storage and processing requirements. In some cases, these penalties might be worth paying as the imagery can always be thinned later on. We found that 80% overlap worked well for our data collection in low canopy tundra environments, in this case all parts of the area surveyed are within 10% of the image centre (near nadir-view for a stabilised sensor) in at least one image and support reliable reconstructions and good quality reflectance map outputs using Pix4Dmapper.

If high amounts of side- and front-lap are not achievable due to limitations of the aircraft or shutter speed of the sensor (*e.g.* due to high flight speeds and wide turns required by fixed-wing aircraft), cross-flight lines can be added to the flight plan (Figure 3a). This will allow the coverage of larger proportions of the surveyed area at near-nadir angles and may reduce BRDF effects. In the case of the Parrot Sequoia, the RGB camera can also be disabled which increases the realised trigger rates for the monochromatic multiband imagery. If problems occur with reconstruction of uniform vegetated surfaces or because of complicated terrains, two diagonal cross-flight lines may be added to the flight plan (Figure 3b), this provides additional coverage of the area and may result in improved reconstructions.

**Figure 3:**
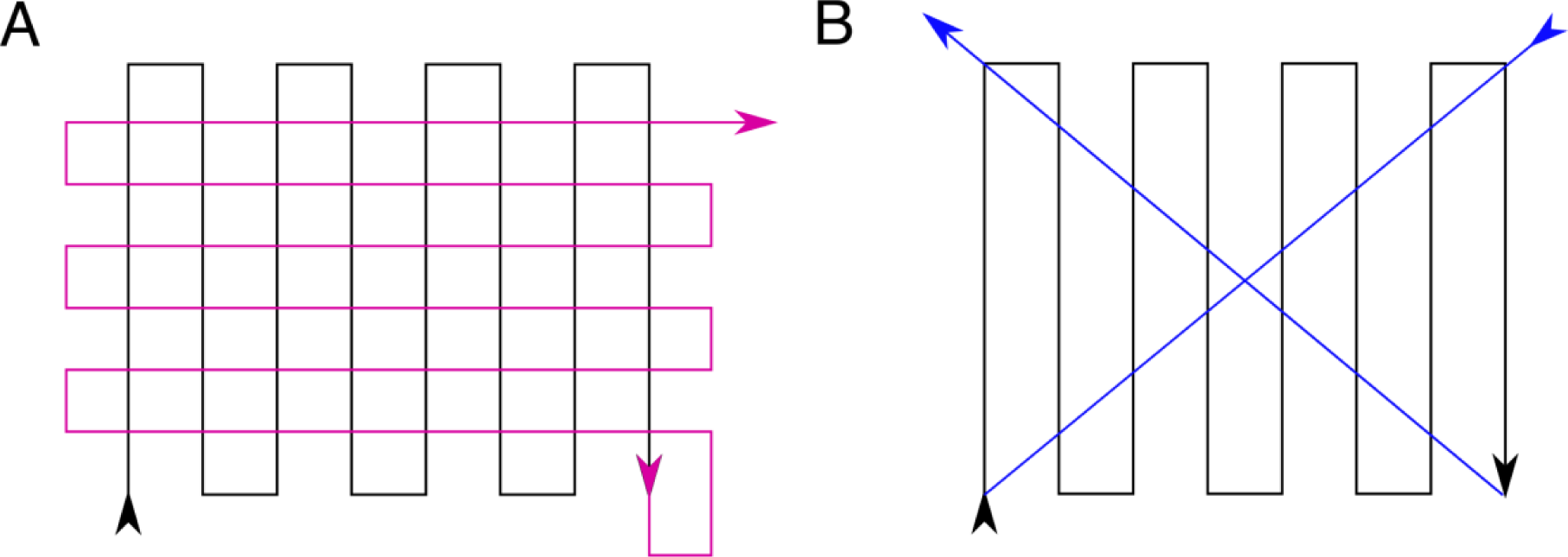
(a) Lawn-mower flight pattern (black) with perpendicular flight lines (pink) to achieve higher overlap and reduce BRDF effects when overlap is limited by aircraft or sensor triggering speed, and (b) Lawn-mover pattern flight path (black) with additional diagonal flight lines (blue) that may aid reconstruction.

The ground sampling distance has a strong influence on the signal to noise ratio. GSD is a function of flight altitude, sensor resolution and optics. Imagery of vegetated surfaces at very small GSDs may contain a lot of noise due to non-uniform reflectance functions and movement of plant parts, such as leaves, between image acquisitions. High amounts of noise hamper key-point matching during SfM-MVS model reconstructions and can reduce the quality of reflectance map outputs, resulting in artefacts, blurry patches and distorted geometry. Pix4D recommends a GSD of 10 cm or more for densely vegetated areas (Pix4D 2018a). Nonetheless, we obtained consistently good results with slightly finer (5 cm) and coarser (15 cm) GSDs for the tussock sedge and shrub tundra vegetation types at our field site QHI in Canada during the data collection campaigns in 2016 and 2017.

When selecting a GSD it is particularly important to consider the scientific objectives of the survey and factor in the scale at which reflectance varies across the area of interest: If the objective is to monitor the distribution of large shrubs, then a larger GSD might be sufficient with the added benefits of reduced noise, the potential to cover larger areas due to higher flight altitudes, less required data storage and faster processing times. In contrast, if the objective is to monitor distribution of small grass tussocks, a smaller GSD might be required with potential penalties due to increased noise in the imagery and reduction in area that can be covered.

### Weather and Sun (Section 4)

Weather and sun are additional factors that influence drone-captured multispectral imagery quality. Most drones will be unable to operate in high winds and rain; but cloud cover and solar position also influence the spectral composition of the ambient light and shadows, thus affecting image acquisition with multispectral drone sensors (Salamí et al. 2014, Pádua et al. 2017). Variation in solar angle may introduce variation in VI estimates even within a single day or flight period (Figure 4). Radiometric calibration of the imagery (Section 6) is a key tool to account for the majority of this variation, but additional steps during flight planning and in-field data collection can be taken to control for some of these factors.

**Figure 4:**
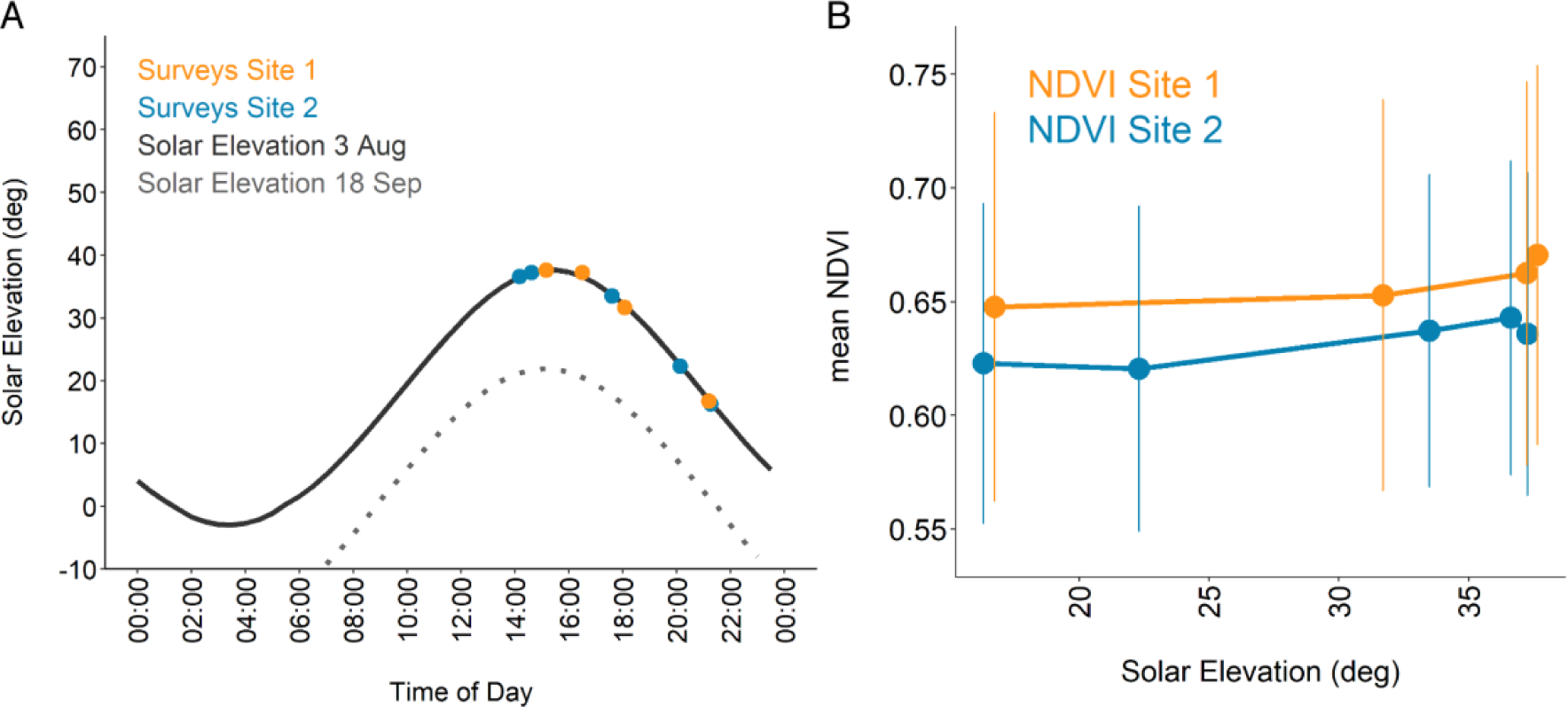
Effect of diurnal solar variation on measured landscape scale mean NDVI. A) Time of day vs. solar elevation for Qikiqtaruk – Herschel Island on 3^rd^ of August 2016 with time-points of repeat surveys shown in B. Light-grey dashed line shows the solar elevation curve for the 18^th^ September 2016, illustrating similar magnitudes of seasonal and diurnal variation across the season at high latitude studies sites such as Qikiqtaruk. B) Effect of solar elevation on mean NDVI for repeat flights of sites on the 3^rd^ of August 2016 on Qikiqtaruk – Herschel Island, highlighting the impact of solar angle and clouds on the mean NDVI values despite radiometric calibration in Pix4D mapper. Bars represent the standard deviation from the mean NDVI (5 cm GSD), illustrating within-site variation at the two 1-ha sites. Absolute differences between highest and lowest solar elevation are just above 0.02 NDVI. Thin stratus cloud cover for all flights except for the flight closest to peak solar elevation (37.22°) at site 2, with low dense cloud, potentially explaining its outlier character.

To minimise variations in solar angle, flights should be conducted as close to solar noon as possible. As a rule of thumb, we recommend a maximum of 2-3 hours before and after solar noon. Seasonal and diurnal variation in solar angle and position can be calculated using solar calculators (such as https://www.esrl.noaa.gov/gmd/grad/solcalc/index.html). At high latitude sites, solar angle will vary across the year in more dramatic ways than at lower latitudes, whereas lower latitudes experience stronger variation in diurnal angle. On clear days, solar position also determines the size and direction of shadows cast on the landscape by micro- and macro-variation in topography (i.e. furrows and ridges, vegetation and hills) (Figure 5).

**Figure 5:**
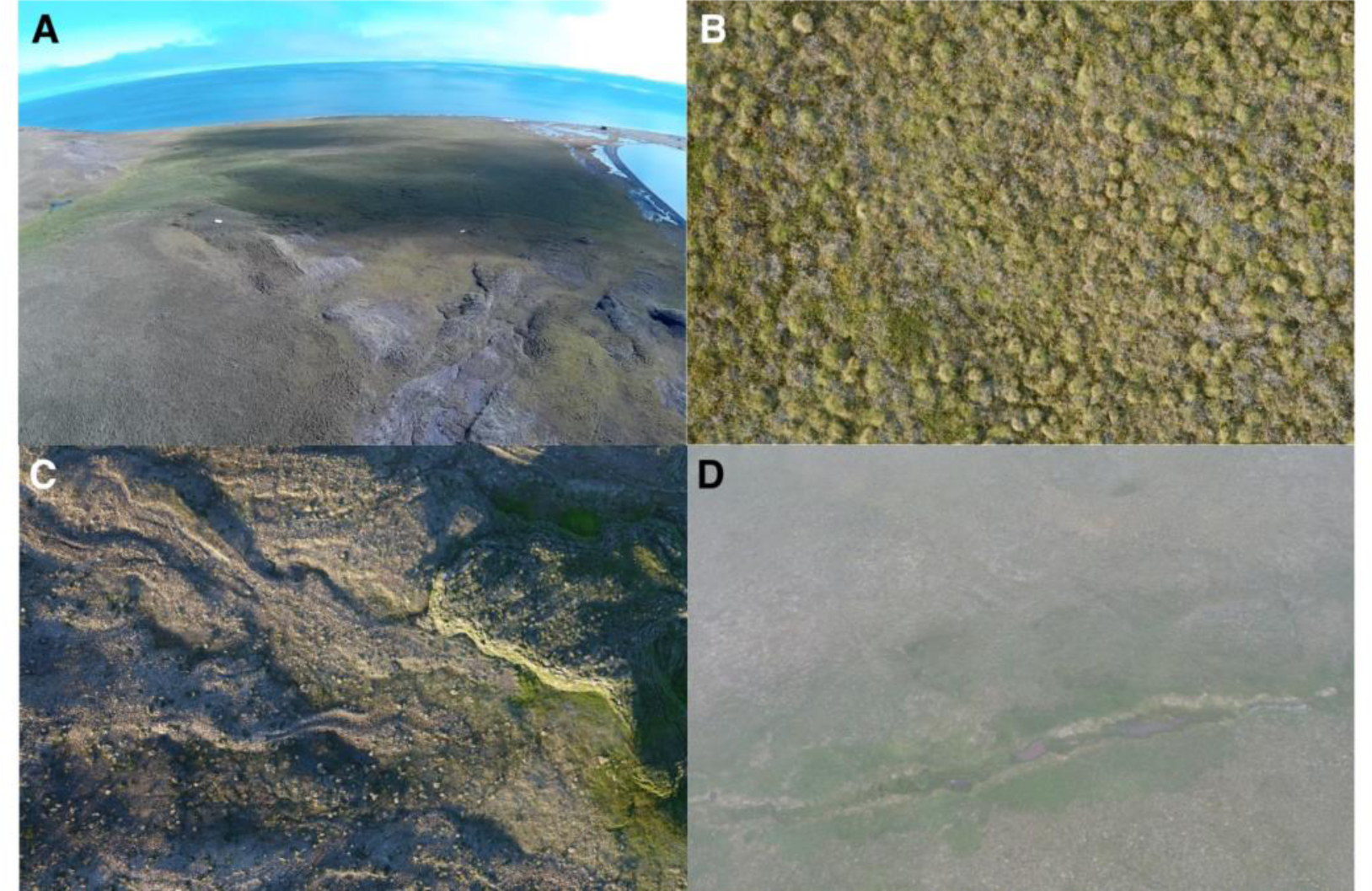
RGB photographs of different cloud and sun angle conditions and their effect on scene illumination. A) “Popcorn” clouds casting well delimitated shadows across the landscape. B) Thin continuous stratus scattering light, resulting in even illumination of the scene and reduced shadows. C) Low solar angle interacting with microtopography, casting shadows across the landscape. D) Fog blurring the imagery and causing uneven illumination.

We recommend recording sky conditions during the flight (Table 2) to account for cloud-induced changes in the spectral composition of light and avoiding days where scattered cumulus clouds (“popcorn-clouds”) are partially shading survey area(s) (Figure 5). The collection of additional meteorological observations such as wind speed (may impact movement of vegetation), temperature and presence of dew/snow may be helpful to account for additional sources of variation in surface reflectance estimates.

**Table 2:** Sky-Codes for qualitative classification of cloud related ambient light conditions. Table courtesy of NERC Field Spectroscopy Facility, Edinburgh UK (2018) based on work by Milton et al. (2009). See also WMO Cloud Identification Guide (World Meteorological Association 2017).

| Sky-Code | Condition |
| --- | --- |
| 0 | Clear sky |
| 1 | Haze |
| 2 | Thin cirrus – sun not obscured |
| 3 | Thin cirrus – sun obscured |
| 4 | Scattered cumulus – sun not obscured |
| 5 | Cumulus over most of sky – sun not obscured |
| 6 | Cumulus – sun obscured |
| 7 | Complete cumulus cover |
| 8 | Stratus – sun obscured |
| 9 | Drizzle |

### Geolocation and Ground Control Points (Section 5)

Accurate geolocation is essential when the image data is: part of a time-series, combined with other sources of geo-referenced data such as satellite or ground-based observations, or used to build structural models. Photogrammetry software packages commonly use two sources of geolocation information: the coordinates of the of the camera during each image capture recorded by the sensor or drone, and/or coordinates of ground control points (GCPs) identified in the imagery. Two problems complicate the accurate geolocation of multispectral imagery products: 1) The accuracy of image geo-tags may be insufficient (ca. ± 2-3 m horizontally) for some scientific applications, and 2) GCPs may be difficult to identify in the low resolution monochromatic images.

The accuracy of geo-tags is limited by the low precision of common drone / sensor GNSS modules. Unless expensive differential positioning systems can be deployed for high accuracy direct georeferencing, the only solution to obtain sub-meter geo-located reflectance maps is to incorporate GCPs whose location is determined in-field with a high accuracy survey grade GNSS. When mapping with the Parrot Sequoia, we recommend the use of a minimum of three to four GCPs well distributed across the area of interest. More may be required for large sites (>1 ha) or sites with varying topography, but higher numbers might not substantially improve 2D geolocation if the area of interest is small and comparatively flat (Figure 6). Additional GCPs not included in geo-rectifying the imagery should be used to assess the accuracy of each reconstruction (Step 10), we recommend at least one additional GCP for this purpose.

**Figure 6:**
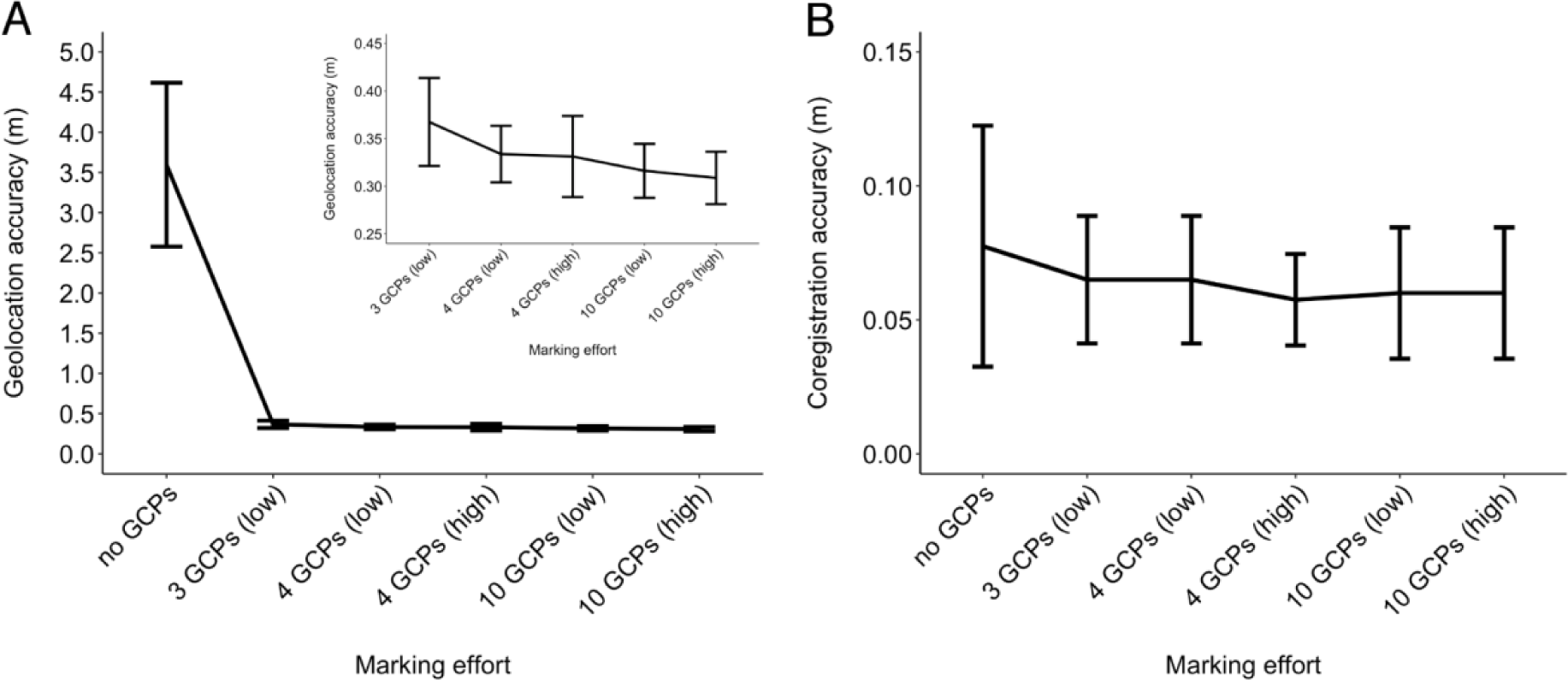
A) Ground Control Point (GCP) marker placement effort and mean geolocation accuracy for eight reflectance maps (red and near-infrared bands) collected at four sites on Qikiqtaruk – Herschel Island. Insert shows data on finer scale excluding the “no GCPs” data point. Images were captured with a Parrot Sequoia at 5 cm per pixel GSD and processed in Pix4D. Error bars indicate standard deviation of the sites from the grand mean. Marking effort was staggered by incorporating 0, 3, 4 or 10 GCPs and increasing the number of images marked per GCP from low (3 images per GCP) to high (8 images per GCP). The relationship suggests diminishing returns for efforts of more than 3 GCPs, with a potential optimum effort-return ratio for 4 GCPs marked at low effort (accuracy approx. 7x GSD). Sites are 1 ha in size and composed of graminoid dominated tundra on predominantly flat terrain with medium amounts of variation in altitude (max 30 m). GCP locations were determined with a survey grade GNSS with a horizontal accuracy of 0.02 m. GCP marker dimensions were 0.265 m x 0.265 m (ca. 5 × 5 GSD) and made from soft plastic or plastic fibres with a black and white triangular sand-dial pattern. Marker contrast was uneven across the monochromatic imagery, resulting in sometimes difficult to distinguish markers. We estimate marker centres were manually identified to ca. two pixels (0.05-0.10 m). Geolocation accuracy of the reflectance maps was assessed by visually locating centre points of 13 GCPs on the final reflectance map outputs in QGIS (QGIS Development Team 2017), this included all GCPs incorporated in the processing. For each reflectance map, the mean absolute distance between visually estimated and computed position was calculated. B) GCP marker placement effort and mean accuracy of co-registration of red and near-infrared reflectance maps from the four sites as in A). The same methods were employed, except the co-registration accuracy was measured as the mean absolute distance between the visually determined locations of the 13 GCPs. The resulting relationship suggests a benefit of including GCPs, but we found no evidence for an improvement with effort of marker placement beyond three GCPs at this flat tundra site.

The compact size and power requirements limit the spatial resolution of CMOS imaging sensors used in multi-camera rigs such as the Parrot Sequoia. This, combined with the reduced spectral bandwidth, can cause difficulties when identifying GCPs in the monochromatic single-band imagery. To achieve maximum visibility of the GCPs, we suggest using square targets composed of four alternating black and white fields arranged in a checkerboard pattern (Figure 7a) with a side length of 7-10x the GSD. The choice of material is important, as white areas of the targets need to reflect strongly across the whole spectrum of the sensor independently of the angle of view (near-lambertian), while black areas should have a low reflectivity to provide a strong contrast. What appears distinctly black and white to the human eye may have similar reflectance properties in the NIR. In our experience, painted canvas and sailcloth are suitable materials that are affordable, readily available and reasonably light. We also achieved good results success with vinyl flooring tiles; however, these can be heavy and therefore impractical in remote field conditions. We strongly recommend testing the visibility of the targets using the multispectral sensors prior field deployment.

**Figure 7:**
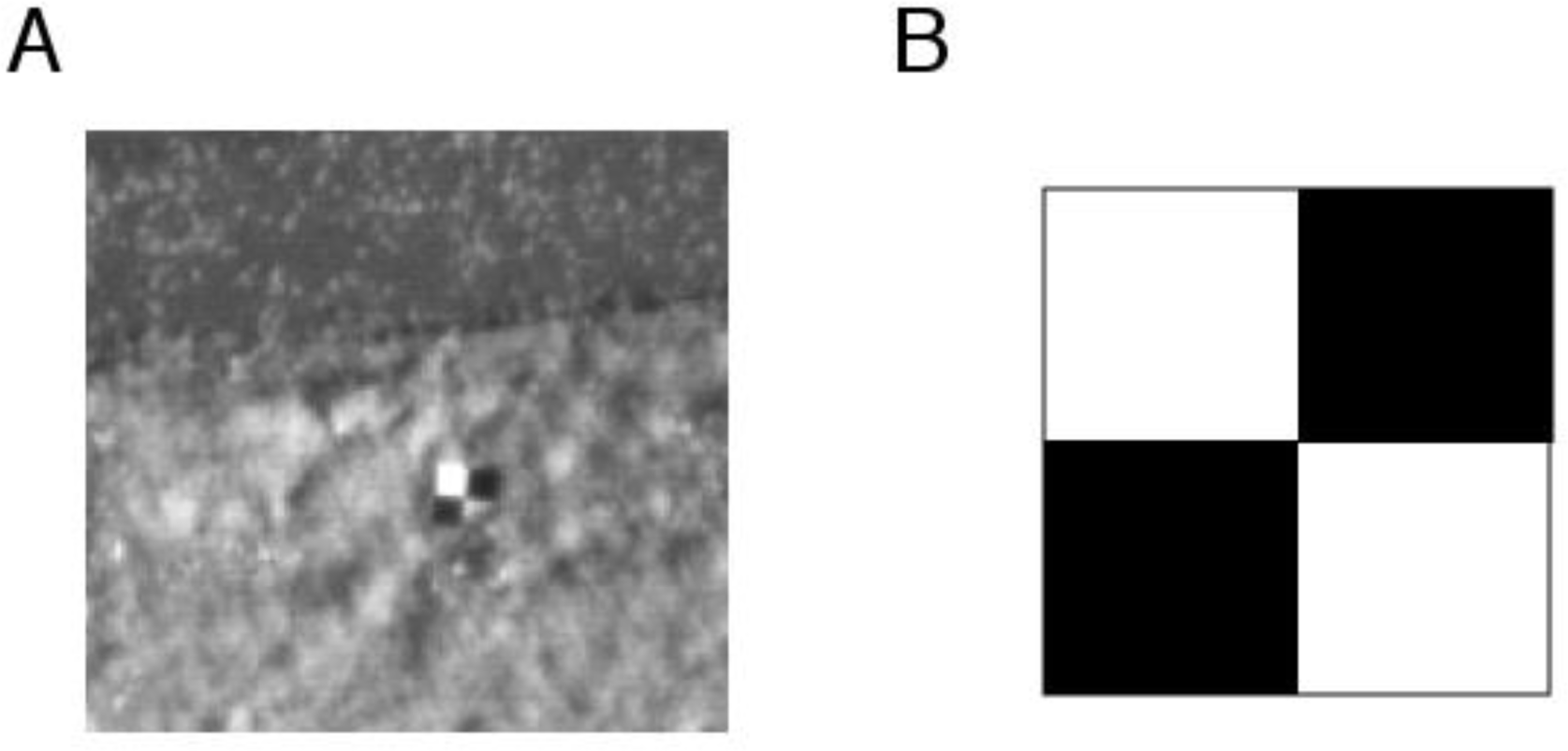
A) Parrot Sequoia near-infrared image of 0.6 m x 0.6 m GCP on grass. This GCP is made from self-adhesive vinyl tiles obtained in a local hardware store. Ground sampling distance: approx. 0.07 m per pixel. Image curtsey of Tom Wade and Charlie Moriarty, The University of Edinburgh. B) Chequerboard pattern suggested for improved visibility of GCP in coarse resolution Parrot Sequoia imagery. Aligning the chequerboard pattern with the sensor orientation can further aid visibility.

Accurate co-registration of pixels among bands is essential when calculating VIs (Turner et al. 2014). Incorporating GCPs in the processing can aid in constraining the relative shifts between the bands. However, we found that increasing the effort in GCP placement in Pix4D for Parrot Sequoia imagery had little impact on constraining the co-registration between bands. High degrees of co-registration (1-2 pixels) were achieved even with the lowest effort of marker placement (Figure 7b). Turner et al. (2014) reported similar levels of co-registration accuracy between reflectance maps of bands collected with a multiband Tetracam mini-MCA (GSD 0.03 m / pixel) at moss sites in Antarctica.

### Radiometric calibration (Section 6)

The aim of the radiometric calibration is to convert at-sensor radiance (in form of DNs) into absolute surface reflectance values, accounting for variation caused by differences in ambient light due to weather and sun, and between sensors types and units (Kelcey and Lucieer 2012). The relationship (empirical line) between image DN values and surface reflectance is established from a sample of pixels covering areas of known reflectance, theoretically this could be a naturally occurring homogeneous area in the area of interest measured with a field spectrometer, but artificial standards (“reflectance targets”) of known reflectance are more commonly used to carry out the calibration.

When processing Parrot Sequoia outputs in Pix4Dmapper a single image is used to calibrate each band (Step 8). A single image is sufficient to establish the empirical line if the sensor response is known and linear (Wang and Myint 2015), as is the case for the Parrot Sequoia (Parrot 2017c). The calibration is carried out by manually selecting the area of the reflectance target on the calibration image (Figure 8) and assigning the known reflectance value of the target. In our experience, a larger samples of pixels produces better calibration results, *i.e.* the more pixels that are taken up by the reflectance target the better. Sample size is likely to be of importance here as it mitigates for variations caused by the inherent noise across the image stemming from the sensor, illumination of the target, and bleeding effects from adjacent non-target surfaces. These findings are consistent with advice from Pix4D (2018b) and MicaSense, who recommend at least 1/3 of the total image size to be covered by the calibration area of the reflectance target (MicaSense 2018b).

**Figure 8:**
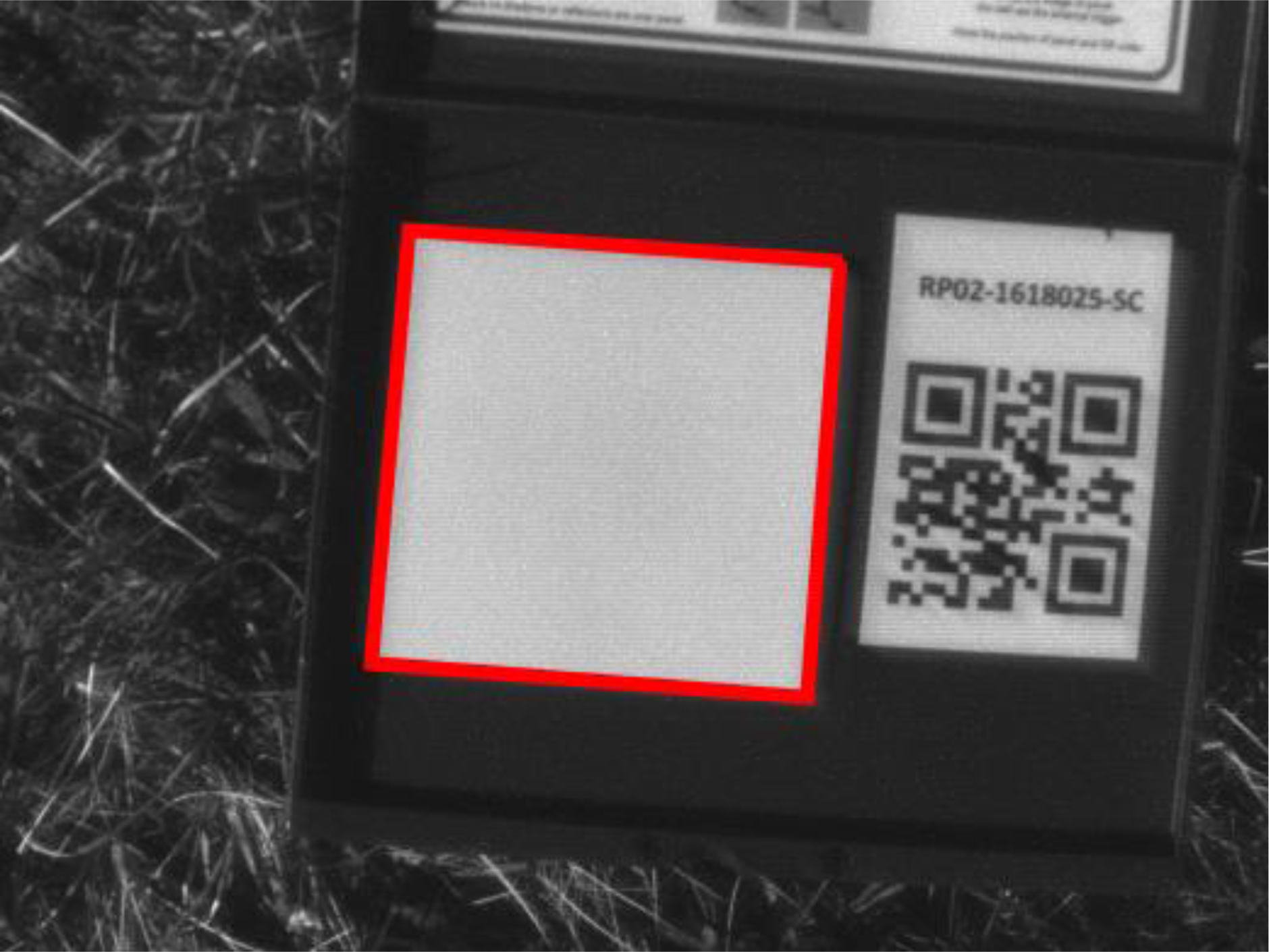
Parrot Sequoia pre-flight radiometric calibration image of a MicaSense Ltd. (Seattle, WA, USA) reflectance target in the near-infrared band. Red box: surface with known reflectance value used for calibration.

Calibration images can be collected either before, after or during the flight. For pre- and post-flight calibration, drone and sensor are held manually above the target and images for all bands are acquired (Step 4). In-flight calibration targets are placed within the area of interest and calibration images acquired during the survey. In-flight targets need to be sufficiently large to ensure a good sample of pixels. Especially when operating in remote areas, weight and size of targets may be limited and quality in-fight calibration imagery can be difficult to obtain. Nonetheless, smaller in-flight reflectance targets (about 100+ pixels = 10+ x 10+ GSD) can be of great use for quality control of the final reflectance map output (see for example Aasen et al. 2015) and may serve as an emergency back-up should pre- /post-flight calibration imagery fail. It is important that both in-flight and pre-/post-flight reflectance targets are placed as level as possible to ensure even illumination of the target surface.

We recommend always obtaining both pre-/post-flight calibration imagery of a reflectance target and, if possible, the use of at least two in-flight reflectance targets for quality control and redundancy. Avoiding overexposure (saturated sensor) and shading of all reflectance targets is critical as this will render the images unusable for radiometric calibration. The Parrot Sequoia has a calibration image acquisition feature for pre-/post-flight calibration accessible via the Wi-Fi interface, which obtains a bracketed exposure and reduces the risk of over-exposure.

When taking pre-/post-flight calibration imagery ensure that as little radiation as possible is reflected onto the target by surrounding objects, including the person taking the calibration picture. Avoiding bright clothing and taking the image with the sun to the photographer’s rear while stepping aside to avoid casting a shadow over the target may reduce the risk of contamination by light scattered from the body (see MicaSense 2018b and, Pix4D 2018b for additional guidance). Aasen and Bolten (2018) observed notable errors introduced to their calibration imagery by the presence and position of the person / drone in the hemisphere above the target, suggesting that the development of reliable calibration methods requires further attention.

It is key that all reflectance targets employed have homogenous and near-lambertian reflectance properties. For pre-/post-flight imagery, we recommend medium sized (approx. 15 × 15 cm) Polytetrafluoroethylene (PTFE) based targets, such as Spectralon (Labsphere 2018), Zenith (Sphereoptics 2018) or similar, due to their durability, off-the shelf calibration and ease of maintenance. Durability and ease of maintenance are particularly important when working in environments with harsh climates. We experienced substantial degradation in commercially manufactured reflectance targets over a single field season (3 months), likely due to exposure to dust, insects, moisture and temperature fluctuations experienced in the Arctic tundra (Figure 9). For larger targets used in-flight we recommend tarpaulins made of canvas, sailcloth, felt or similar materials (see Ahmed et al. 2017, Crusiol et al. 2017, Mosaic Mill Ltd. 2018). Nonetheless, a variety of other materials have also been successfully employed as reflectance targets (Laliberte et al. 2011, Turner et al. 2014, Wang and Myint 2015, Aasen et al. 2015, Wehrhan et al. 2016, Dash et al. 2017).

**Figure 9:**
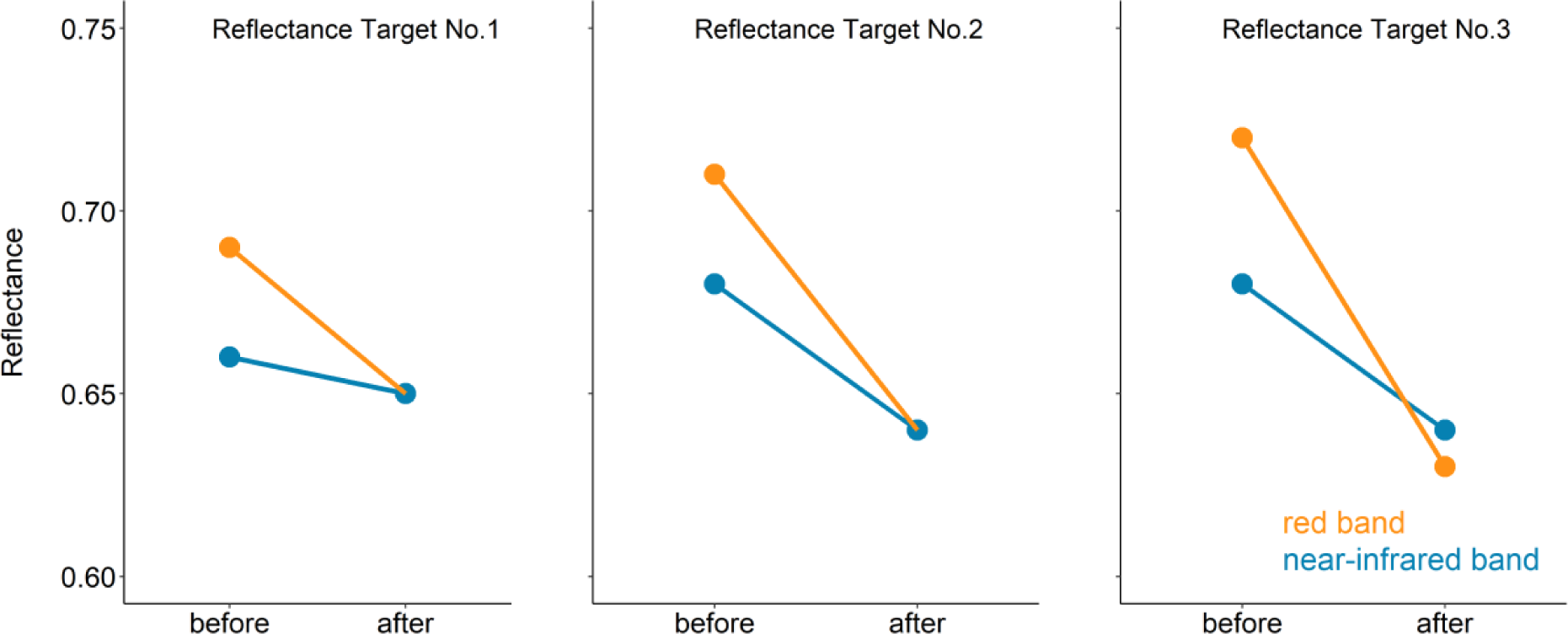
Decrease in reflectance values of three reflectance targets before and after a three-month field season in the Arctic tundra on Qikiqtaruk – Herschel Island. Loss in reflectance is likely due to degradation in the harsh environmental conditions (dust, insect debris, moisture and temperature fluctuations). Across the field seasons in 2016 and 2017 we saw 4-10% reduction in reflectance across targets from different suppliers, composed of different materials.

Target maintenance and quality control is essential (also discussed by Wang and Myint 2015). Changes in target reflectance can have notable effects on the calibration outputs (Figure 10). It is key to handle targets as carefully as possible to avoid surface degradation. We recommend regular cleaning according to manufacturers’ guidance and frequent re-measurement of reflectance values. Field spectroscopy facilities can provide assistance and expertise in obtaining and maintaining targets. Re-measurement of the reflectance values can be carried out in-field prior each flight (e.g. Laliberte et al. 2011). However, this might not always be feasible when operating in remote areas, in which case careful handling and maintenance before and after a field season may have to suffice.

**Figure 10:**
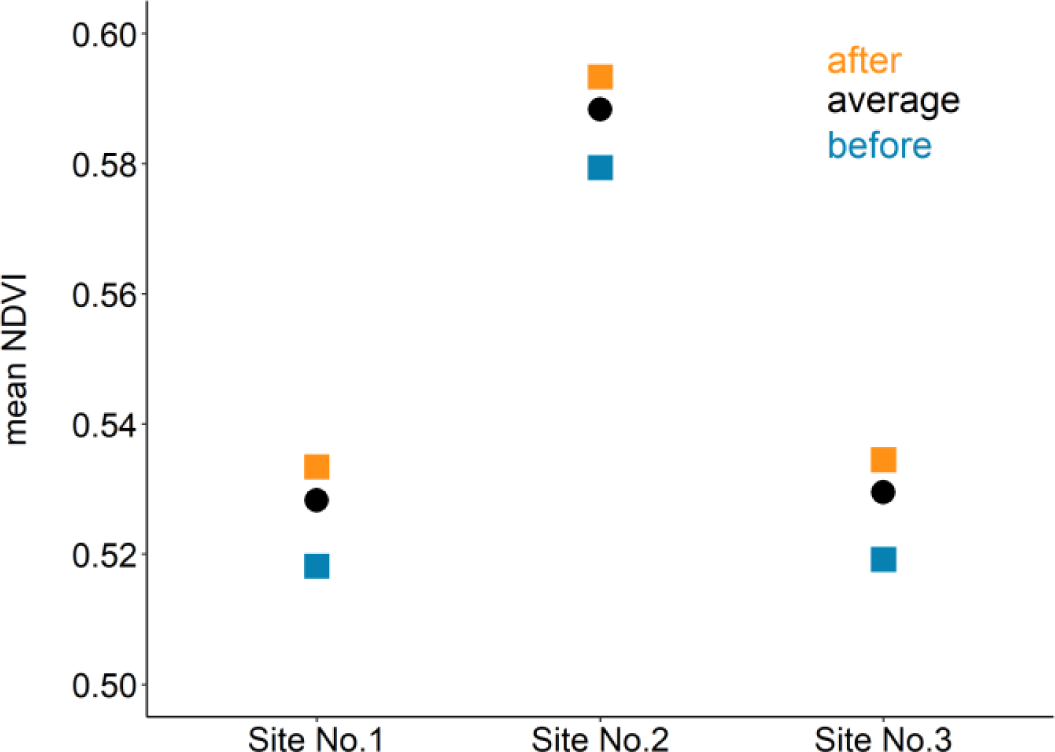
Mean NDVI value for three graminoid tundra sites (1 ha each) on Qikiqtaruk – Herschel Island based on red and near-infrared reflectance maps calibrated with three different reflectance values for the reflectance target No. 1 (Figure 9): before and after degradation, and the average between the two values. Surveys where flown at the beginning of the season when little to no degradation of the target is expected to have occurred. Before and after values differ by about 0.015 in absolute NDVI, suggesting an overestimation of NDVI when after values are used for the early season surveys.

Optical filters directly affect the radiation reaching the sensor and influence the relationship between surface radiance and image DN, see Kelcey and Lucieer (2012) for further discussion. It is therefore essential that all radiometric calibration imagery and survey photographs are consistently taken either with or without the removable filter. The Parrot Sequoia is shipped with a protective lens cover (a clear filter), which can be useful when operating in difficult terrains such as the tundra where rough landings are possible, which could scratch the sensor lenses. Parrot does not characterise the transmissivity of the protective lens covers shipped with the Sequoia. As the presence / absence of filters is difficult to detect *post hoc* during automated processing (such as online cloud services), Parrot recommends refraining from using them during multispectral data acquisition flights (Parrot 2017b).

We measured the transmissivity of the filters shipped with two Sequoias obtained in 2016 (Figure 11). We observed a small reduction in transmitted radiation across all four bands, and a small effect of angle of view across the horizontal field of view on the radiation transmitted in the near-infrared band. These findings suggest that the protective lens cover may be used with little to no effect on the final reflectance map outputs, if the filter is applied consistently for all flights under comparison.

**Figure 11:**
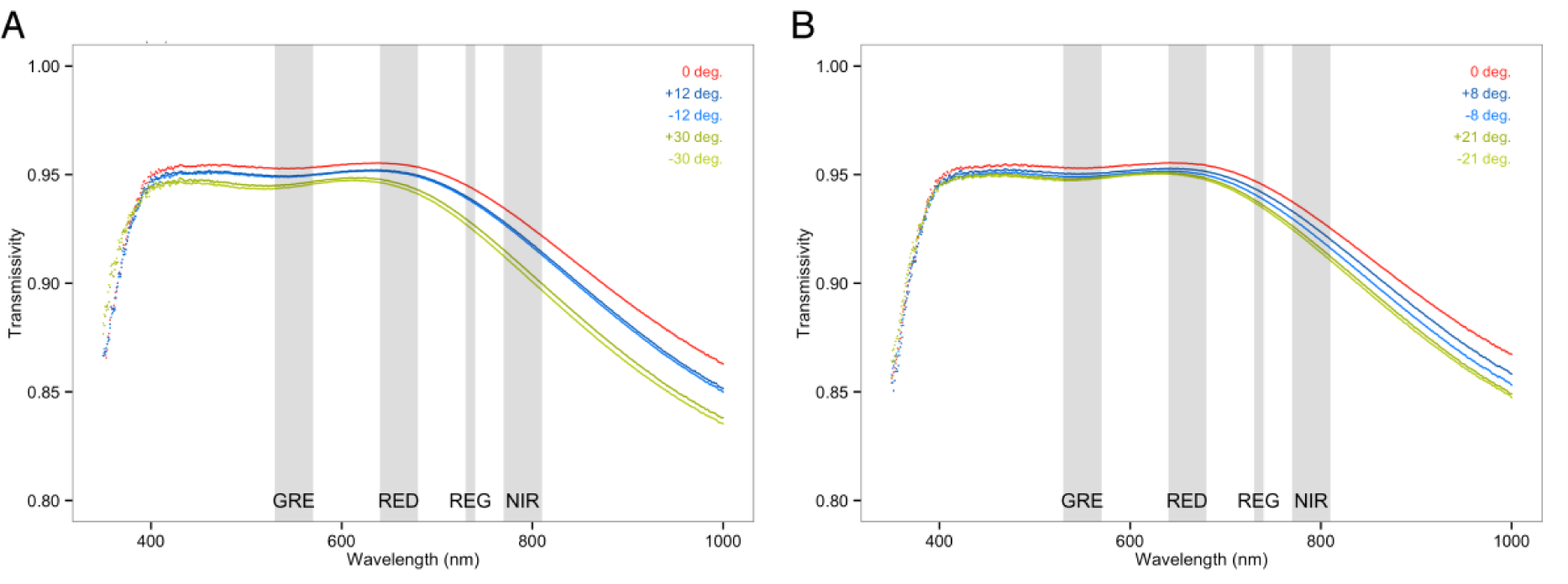
Transmissivity of Parrot Sequoia Lens-Protector filter across the a) horizontal and vertical field-of-view of the Sequoia Sensor. The overall small reductions in transmitted light and the small effect of angle across field-of-view suggest that little to no impact on reflectance map outputs acquired with the filter can be expected.

### Estimated combined error

We estimate that the combined effect of the main sources of error discussed in this manuscript – if not properly accounted for - could be as much as 0.094 in magnitude for landscape level estimates (1 ha mean) in NDVI for the drone surveys conducted with a Parrot Sequoia at 5 cm GSD at our Arctic research site Qikiqtaruk during the 2016 field campaign (Fig. 13). This combined error equates to approximately 10-13% of the peak growing season NDIV (0.60 - 0.68) of the tussock-sedge and dryas-vetch tundra types at the site. These estimates highlight the importance of controlling for these sources of error, by carrying out radiometric calibration, surveying at constant solar angles, monitoring reflectance target degradation and using the protective lens cover consistently. Nonetheless, a notable error will remain even if everything except cloud conditions is controlled for, we estimate that our ability to then confidently detect change in landscape scale (1 ha) mean NDVI is limited to differences above 0.02 - 0.03 in absolute magnitude across space and time.

**Figure 12:**
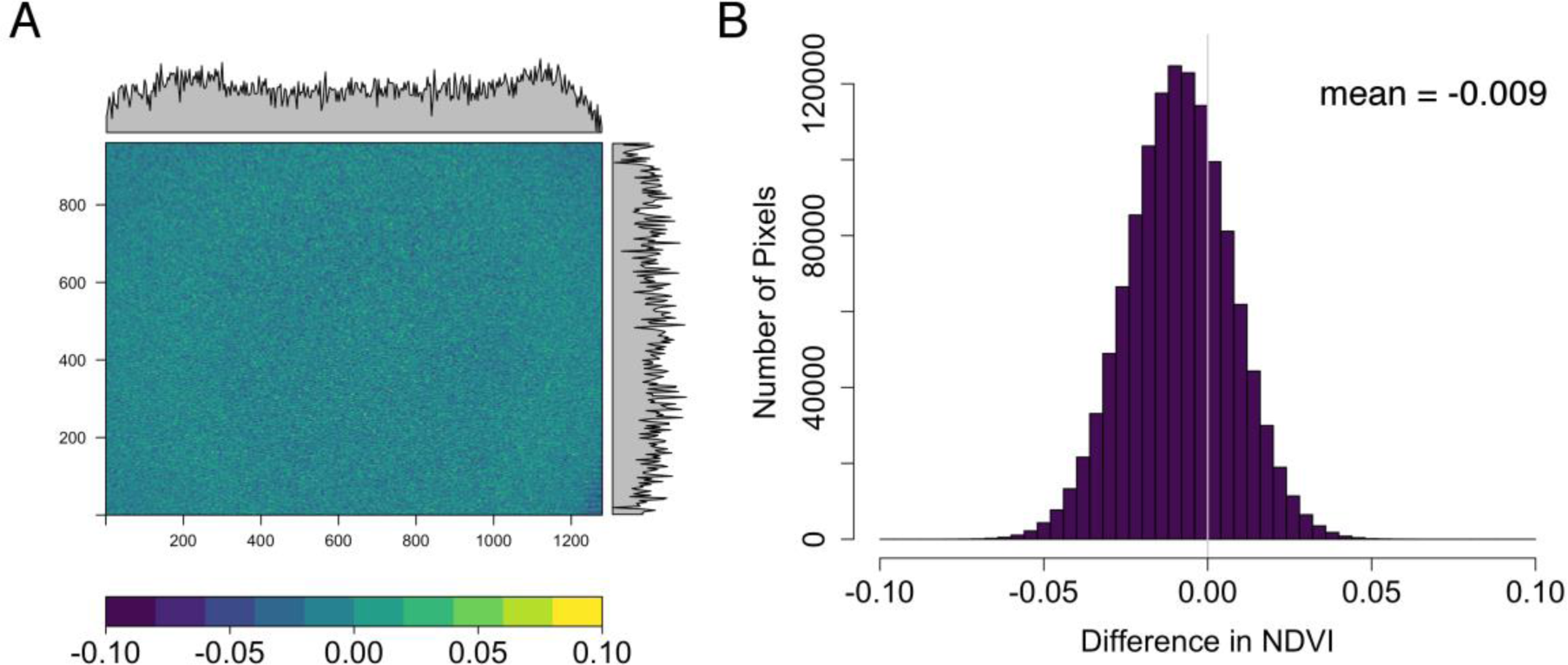
Raster plot (A) and histogram (B) of pixel by pixel differences in NDVI values of a homogenously illuminated integrating sphere with and without the Parrot Sequoia protective lens cover. Margins in the raster plot show mean differences for the pixel columns and rows respectively.

**Figure 13:**
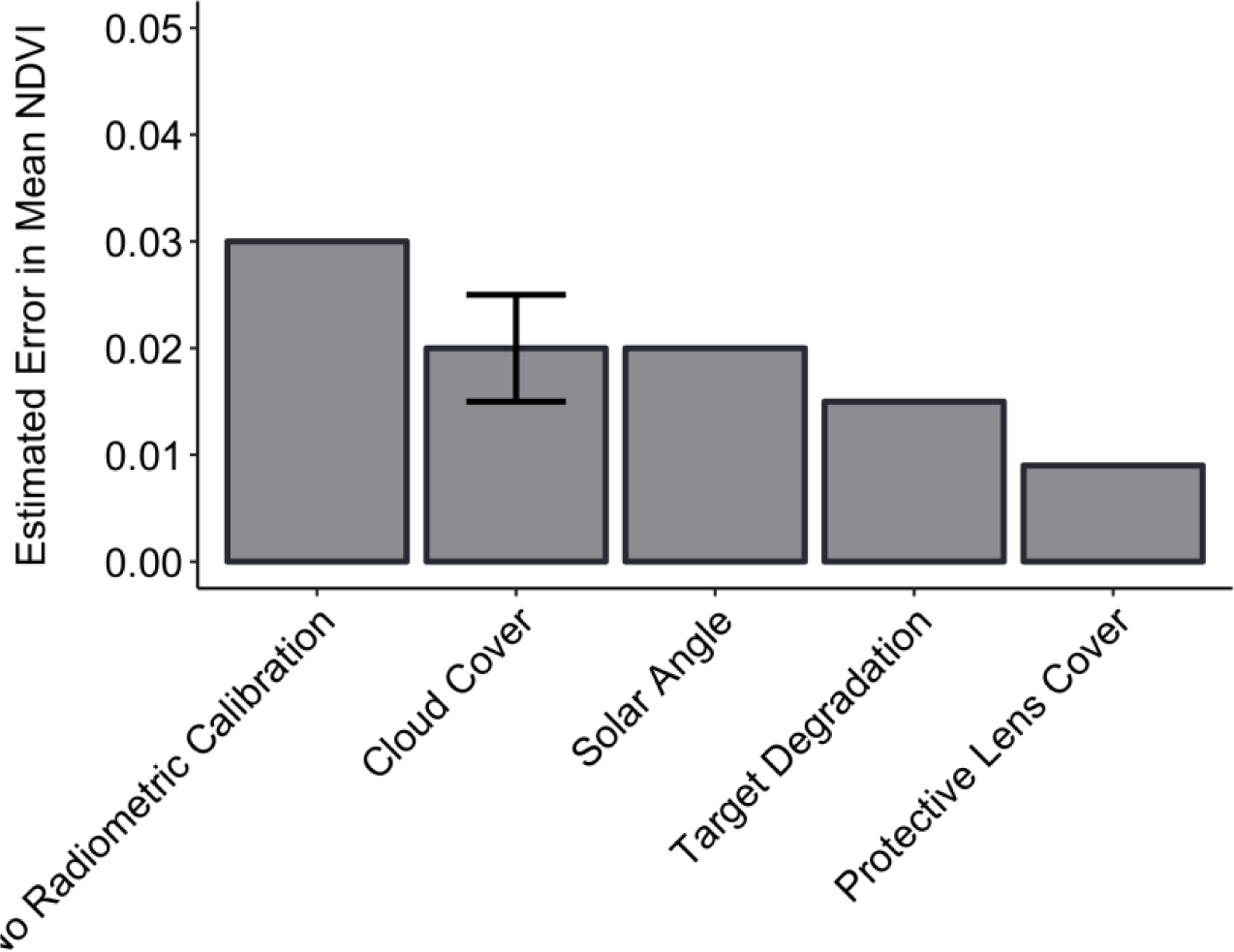
Estimated effects of the five main sources of errors discussed in this manuscript on the mean NDVI of 1 ha tundra plots on Qikiqtaruk surveyed in 2016 with a Parrot Sequoia at 50m flight altitude (5 cm GSD). The five sources of error are: 1) The estimated average deviation from the calibrated mean NDVI compared to a survey without radiometric calibration carried out. 2) The deviation in estimated mean NDVI when comparing clear sky to continuous cloud cover conditions (lower error bar: thick stratus, upper error bar: thick cumulus) even if radiometric calibration is carried out. 3) The estimated deviation of mean NDVI caused by changes in solar elevation from solar noon to evening during peak growing season at our field site in the Arctic (about 20° drop – roughly equivalent to the difference between start/end and mid growing season) even if radiometric calibration is carried out. 4) The estimated effect of target degradation on mean NDVI across a three-month field season. 5) The error introduced by the protective lens cover if used and removed inconsistently between flights in comparison. These estimates are based on both data presented in this manuscript and manuscripts in preparation. We would like to urge caution when transferring these estimates to other sensors / set ups and ecological systems. The estimates are presented here with the purpose of giving the reader a feel for the relative importance of the sources of error discussed in this manuscript.

## Conclusions

Vegetation monitoring using drones could provide key datasets to quantify vegetation responses to global change (Anderson and Gaston 2013, Salamí et al. 2014, Torresan et al. 2017). However, accurately quantifying and accounting for the common sources of error and variation is a key part of the workflow for a drone ecologist (Aasen et al. 2015, Manfreda et al. 2018). As technologies advance and our understanding of multispectral drone products increases we may be able to better quantify the sources of error and improve our measures to account for them; however, it is critical that the drone data collection of today is done as cautiously and rigorously as possible as it will provide the baseline for future ecological monitoring studies.

The rapid and ongoing development of drone and sensor technology (Anderson and Gaston 2013, Pádua et al. 2017) has made the collection of multispectral imagery with drones accessible to many ecological research projects, even those operating with small budgets. Despite the plug-and-play nature of the latest generation of multispectral sensors, such as the Parrot Sequoia and the MicaSense RedEdge, a handful of factors require careful consideration if the aim is to collect high quality multispectral data that is comparable across sensors, space and time. For example, variation in ambient light and sensors require radiometric calibration of the imagery, and ground control points may be necessary to achieve accurate geolocation of reflectance and vegetation index maps (Kelcey and Lucieer 2012, Turner et al. 2014, Salamí et al. 2014, Aasen et al. 2015, Pádua et al. 2017).

In this manuscript, we suggested a standardized workflow for multispectral drone surveys, discussed the technical aspects and challenges of multispectral drone sensors, flight planning, the influence of weather and sun, as well as aspects of geolocation and radiometric calibration. We believe that these key factors, if properly accounted for, will allow for the majority of multispectral drone surveys to produce data that is comparable across different study regions, plots, sensors and time. We encourage ecologists and other researchers to incorporate these methods and perspectives in their planning and data collection to promote higher data quality and allow for cross site comparisons. Standardised procedures and practises across research groups (e.g., those developed by the HiLDEN network) have the potential to provide highly-valuable baseline data that can be used to address urgent and emerging topics, such as identifying the landscape patterns and processes of vegetation change.

## Acknowledgements

Much of this manuscript would have not been possible without the valuable input from Chris MacLellan and Andrew Gray at the NERC Field Spectroscopy Facility at the Grant Institute in Edinburgh. We would also like to thank Andrew Gray for providing feedback on an earlier version of this manuscript and Tom Wade from the University of Edinburgh Airborne GeoSciences Facility University of Edinburgh Airborne GeoScience facility for his ongoing support of our drone-based endeavours in the Arctic.

Funding for this research was provided by NERC through the ShrubTundra standard grant (NE/M016323/1), a NERC E3 Doctoral Training Partnership PhD studentship for Jakob Assmann (NE/L002558/1), a research grant from the National Geographic Society (CP-061R-17) and a Parrot Climate Innovation Grant for Jeffrey Kerby, a NERC support case for use of the NERC Field Spectroscopy Facility (738.1115), equipment loans from the University of Edinburgh Airborne GeoSciences Facility and the NERC Geophysical Equipment Facility (GEF 1063 and 1069).

